# Hapo-G, Haplotype-Aware Polishing Of Genome Assemblies

**DOI:** 10.1101/2020.12.14.422624

**Authors:** Jean-Marc Aury, Benjamin Istace

## Abstract

Single-molecule sequencing technologies have recently been commercialized by Pacific Biosciences and Oxford Nanopore with the promise of sequencing long DNA fragments (kilobases to megabases order) and then, using efficient algorithms, provide high quality assemblies in terms of contiguity and completeness of repetitive regions. However, the error rate of long-read technologies is higher than that of short-read technologies. This has a direct consequence on the base quality of genome assemblies, particularly in coding regions where sequencing errors can disrupt the coding frame of genes. In the case of diploid genomes, the consensus of a given gene can be a mixture between the two haplotypes and can lead to premature stop codons. Several methods have been developed to polish genome assemblies using short reads and generally, they inspect the nucleotide one by one, and provide a correction for each nucleotide of the input assembly. As a result, these algorithms are not able to properly process diploid genomes and they typically switch from one haplotype to another. Herein we proposed Hapo-G (Haplotype-Aware Polishing Of Genomes), a new algorithm capable of incorporating phasing information from short reads to polish genome assemblies and in particular assemblies of diploid and heterozygous genomes.

## INTRODUCTION

Long-read technologies commercialized by Pacific Biosciences (PACBIO) and Oxford Nanopore Technologies (ONT) are able to sequence long DNA molecules but at the cost of a higher error rate at least for standard protocols. Their throughputs are sufficient to generate complex genomes(1–5) and their costs are almost compatible with their use in large-scale resequencing projects(6–8). Standard genome assemblies currently rely on a combination of several technologies, making it possible to generate complete assemblies in terms of both repetitive and coding regions. The quality of the consensus relies heavily on the use of short reads and the choice of a polishing algorithm.

One of the most popular polishing algorithms, Pilon(9), was developed several years ago, before the advent of the long-read era and was originally designed to detect variants and improve microbial genome assemblies. With the increasing popularity of long-read technologies, public databases now contain a large collection of very contiguous assemblies, but even if the overall reported quality seems sufficient, local errors can critically affect protein prediction(10). Aware of this issue, the bioinformatic community has developed several tools over the past two years(11–15). Most of the tools (Pilon, Racon, NextPolish, HyPo and POLCA) are based on short-read alignment, ntEdit is the only method that uses a kmer approach and NextPolish combines both strategies (Table 1). After aligning the short reads, the algorithms detect errors by examining the pileup of bases from the reads (Pilon, NextPolish), by generating a consensus using Partial Order Alignment (Racon, HyPo) or by detecting variants (POLCA). While these tools are capable of correcting most of the errors in a draft assembly generated using long reads, we have observed frequent issues when correcting heterozygous regions. Indeed, the case of diploid genomes is particularly problematic since in this case the long-read assembly is composed of collapsed homozygous regions and duplicated allelic regions which will complicate the correct alignment of short reads. As the existing tools work locally and not at the scale of a 150bp read and its mate, they frequently generate a mixture of haplotypes. Switching between haplotypes is problematic for the alignment of short reads and variant calling, but it can also affect the coding sequence of genes. As an example, when we were dealing with a long-read genome assembly, we observed that the pilon correction was not able to restore a deletion in a heterozygous coding region (Figure 1). This simple observation motivated our need to develop a new polishing algorithm.

**Table 1.** General characteristics of existing polishing algorithms. These seven tools were evaluated in our benchmark with the specified parameters.

|  | Reference | Publication Date | Version | Read alignment | Integrated aligner | Parameters used in the benchmark |
| --- | --- | --- | --- | --- | --- | --- |
| Hapo-G | This study | - | 0.1 | yes | BWA | -t 36 |
| POLCA | Ziminn AV. et al.(14) | 2020 | 3.4.2 | yes | BWA | -t 36 |
| HyPo | Kundu R. et al. (13) | 2019 | 1.0.3 | yes | no | -s size*<br>-c 180<br>-t 36 |
| NextPolish | Hu J. et al.(12) | 2019 | 1.3.2 | yes | BWA | task = best<br>parallel_jobs = 6<br>multithread_jobs = 6<br>genome_size = auto |
| ntEdit | Warren RL. et al. (11) | 2019 | 1.3.2 | no | NA | -k 40 -t 36<br>--outbloom<br>--solid<br>ntEdit : -m 1 -t 36 |
| Racon | Vaser R. et al. (15) | 2017 | 1.4.3 | yes | no | -t 36 |
| Pilon | Walker BJ. et al. (9) | 2014 | 1.23 | yes | no | --threads 36 |

\* size= 120Mb (*Arabidopsis thaliana*), 120Mb (*Solanum tuberosum L*), 100K (synthetic sequence)

**Figure 1.**
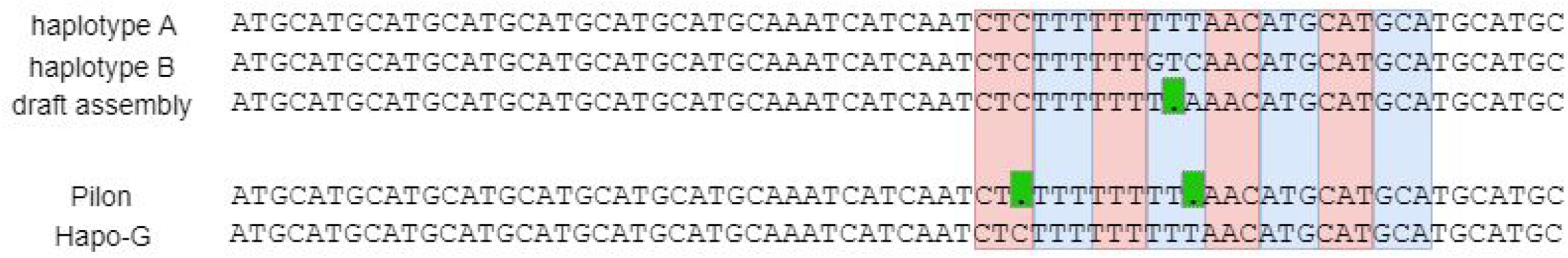
Example of a deletion in a coding frame. The two haplotypes have consistent coding frames (codons are alternatively colored in red and blue) and the draft assembly contains a deletion in a stretch of T’s (green box). Pilon was not able to restore the coding frame and add a second frameshift. In comparison Hapo-G was able to restore haplotype A.

Here we present Hapo-G (pronounced as apogee), a new method dedicated to the polishing of genome assemblies. This algorithm tends to phase the assembly while correcting the sequencing errors. We compare Hapo-G with existing short-read polishers (HyPo, NextPolish, Pilon, POLCA and Racon) and show that Hapo-G is not only comparable to existing methods for polishing draft assemblies, but is also faster and tends to decrease jumps between haplotypes. Hapo-G is written in C, uses the hts library(16) and is freely available at http://www.genoscope.cns.fr/hapog.

## MATERIAL AND METHODS

### Hapo-G algorithm

Hapo-G, like most existing tools, requires a sorted bam file containing the short-read alignments on the draft genome. These short-read alignments could have been generated using bwa mem(17), minimap2(18) or any other alignment tool capable of producing a bam file. Hapo-G maintains two stacks of alignments, the first (all-ali) contains all the alignments that overlap the currently inspected base, and the second (hap-ali) contains only the read alignments that agree with the last selected haplotype. Hapo-G selects a reference alignment and tries to use it as long as possible to polish the region where it aligns, which will minimize mixing between haplotypes (Figure 2).

**Figure 2.**
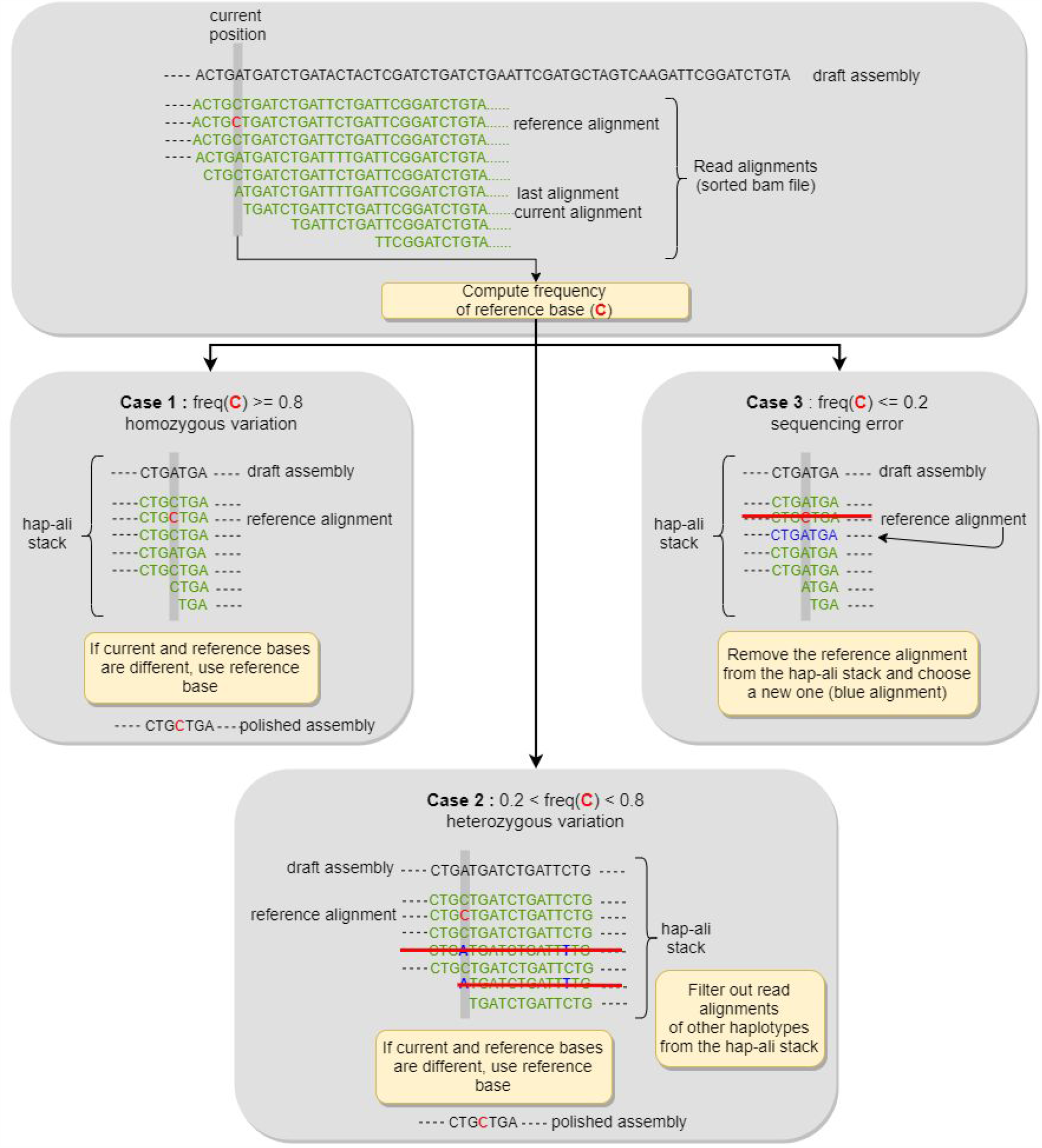
Description of the Hapo-G algorithm. Two stacks of alignments are stored, all-ali which contains all the alignment of a specific region (not shown) and hap-ali which contains reads from the same haplotype. The bam file is processed iteratively, and for each input alignment, Hapo-G will polish the region (draft genome is in black) between the start of the last alignment and the start of the current alignment. The reference alignment is the one used as the backbone for error-correction. Once the frequency of the reference base (in red) is computed, the position is classified as homozygous (case 1, left panel), heterozygous (case 2, lower panel), or sequencing error (case 3, right panel).

**Figure 3.**
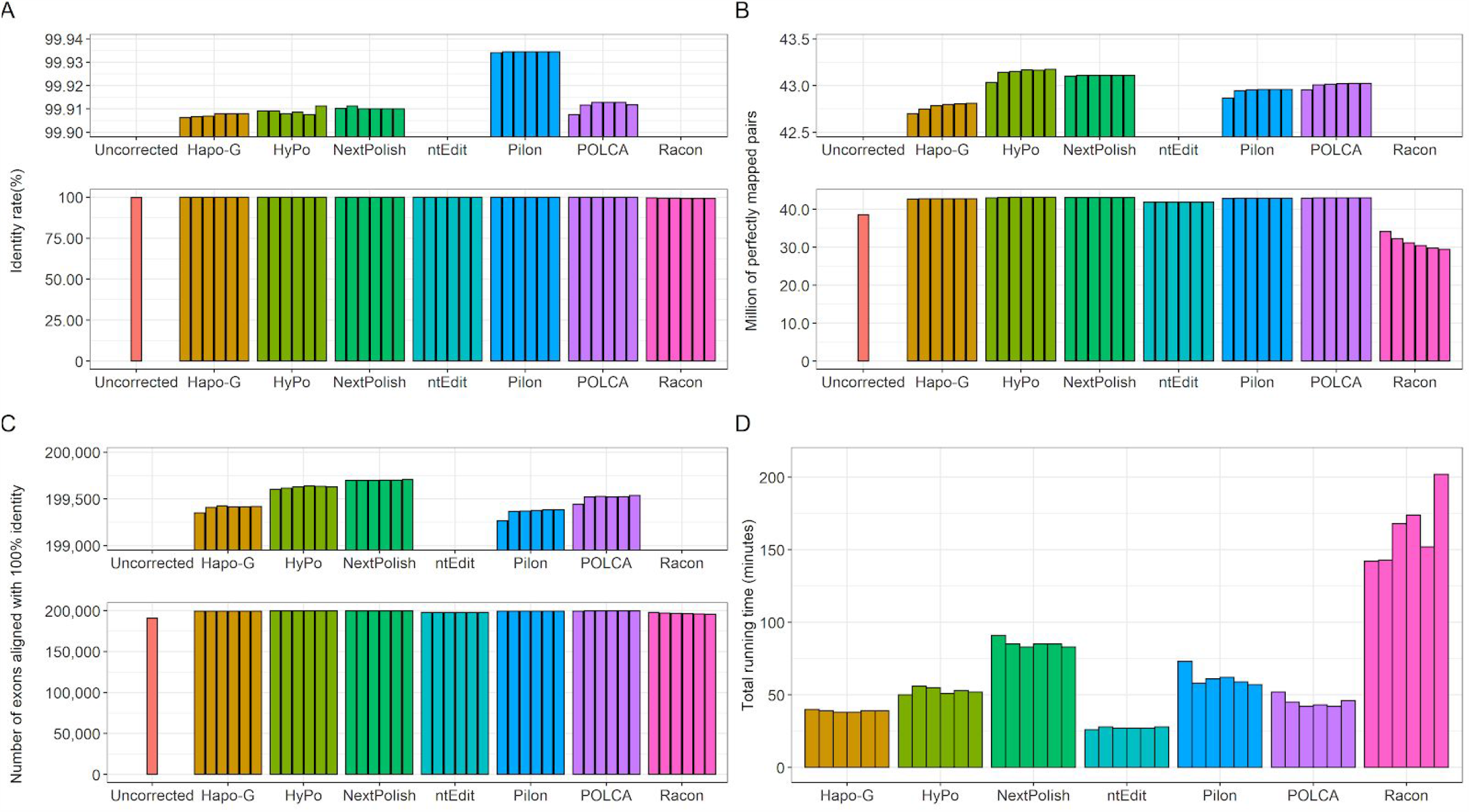
Comparison of polishing algorithms on the *Arabidopsis thaliana* genome assembly. Lower panels of A, B and C show the full distribution and the upper panels are a zoom on the higher values. **A**. Identity rate of assemblies after each round of polishing, when compared to the *Arabidopsis thaliana* reference genome. **B**. Number of Illumina pairs mapped perfectly on each assembly. **C**. Number of Arabidopsis thaliana exons aligned with 100% identity after each round of polishing. **D**. Run times of polishing tools, for each polishing round.

Hapo-G performs the polishing sequentially and scans the input bam file of alignments sorted by position. For each input alignment (called current alignment), Hapo-G polishes each nucleotide in the region between the last recorded position and the start position of the current alignment (called current position), after which the current alignment will be added to both stacks (Figure 2).

### Polishing of a given nucleotide in the draft assembly

First, the two stacks (all-ali and hap-ali), if they are not empty, are cleaned to remove any alignment that does not overlap with the current position. In case the reference alignment has been deleted, a new alignment is selected from the hap-ali stack (the read alignment that ends closest to the current position). If the coverage at the current position is below a threshold, set at three reads, the current base in the draft sequence remains unchanged. Otherwise, the nucleotide of the reference alignment (called the reference base) is extracted and the frequency of this reference base is calculated in the all-ali and hap-ali stacks. Based on its frequency, the current position is tagged as a homozygous site, a heterozygous site or a sequencing error (Figure 2).

The position is classified as homozygous if the frequency of the reference base is greater than 0.8 and at least 3 reads from the hap-ali stack are in accordance. If the reference base and the nucleotide of the draft assembly (the current base) are different, the current base is replaced by the reference base.

The position is classified as heterozygous if the frequency of the reference base is between 0.2 and 0.8 and the hap-ali stack contains at least six reads. If the reference base and the current base are different, the current base is replaced by the reference base. In addition, any read alignments that do not have the same base as the reference base at the current position will be removed from the hap-ali stack. Indeed, they may represent a second haplotype. Importantly, when a read is removed from the stack, its name and its mate name are added to a hash table. The corresponding read alignments will be ignored when encountered later while polishing the current sequence. The hash table is empty when the end of the current sequence is reached.

### Usage and parallelisation of Hapo-G

The polishing step of Hapo-G is wrapped in a python script which manages the pre and post processing steps. First, the wrapper indexes the genome and maps the short reads on the draft assembly using bwa mem. The polishing step, written in C using the htslib, is not multithreaded but can be easily parallelized by splitting the input fasta file as well as the alignment file. This divide and conquer strategy makes it possible to speed up the polishing step, and allows to take advantage of a wide range of computing architectures.

### Generation of benchmarking datasets

#### Homozygous genome assembly - *Arabidopsis thaliana*

We downloaded the Nanopore data produced by Michigan State University (Table 2) and assembled these data using the Flye assembler(19) (v2.8.1) with the parameter ‘-g 120m’ to indicate a genome size of 120Mb. We obtained an assembly of 118Mb, comprising 16 large contigs and an N50 contig of 14.8Mb (Table 2). This assembly was used as input of the Medaka polisher(20) (v1.2.0), in conjunction with the Nanopore reads and the ‘-m r941_min_high_g303’ parameter was applied, in order to choose a suitable model for this type of data. This assembly was used to compare short-reads polishing tools.

**Table 2.** Datasets and long-read assemblies generated for the benchmark. Coverages were computed using a genome size of 120Mb and 1,600Mb for *Arabidopsis thaliana* and *Solanum tuberosum* respectively.

|  |  | *Arabidopsis thaliana*<br>Col-0 | Synthetic sequence | *Solanum tuberosum* L.<br>RH89-039-16 |
| --- | --- | --- | --- | --- |
| Illumina | Accession number | SRR12136403 | NA | PRJNA573826 |
|  | Read Length (bp) | 2x150 | 2x150 | 2x250 |
|  | Coverage | 176 X | 50 X | 47 X |
| Nanopore | Accession number | SRR12136402 | - | PRJNA573826 |
|  | Reads N50 (bp) | 18,827 | - | 25,280 |
|  | Coverage | 95 X | - | 75 X |
| Assembly | Number of contigs | 238 | 1 | 11,070 |
|  | Cumulative size | 119,992,853 | 102,000 | 1,332,417,447 |
|  | Contig N50 (bp) | 14,841,396 | 102,000 | 440,422 |

#### Synthetic diploid sequence

We generated a 100Kb sequence using an Homo sapiens model(21) and created two haplotypes by incorporating, each time, 100 random mutations into the initial 100Kb sequence(22). We added two random 1Kb sequences to both ends of the two haplotypes to avoid mapping issues on first and last nucleotides. Illumina short-reads were generated from both haplotypes using ART(23) software (version 2.5.1) and the following parameters: -ss HSXt -p -l 150 -f 25 -m 200 -s 10. From the two haplotype sequences, we generated an haploid sequence by alternatively retaining 60 nucleotides of each sequence and adding 2,000 random mutations(22) to simulate sequencing errors. The resulting haploid sequence is a mixture of the two haplotypes and is used to test the ability of each polisher to correct errors and phase the draft sequence.

#### Heterozygous genome assembly - *Solanum tuberosum L*

We downloaded the Nanopore data produced by the authors of a recent article by Zhou et al.(24) describing the diploid assembly of *Solanum tuberosum* (Table 2). The long-read dataset was assembled using the Flye assembler(19) (v2.8.1) with the parameter ‘-g 1600m’ to indicate a genome size of 1600Mb (representing the length of the diploid genome). The resulting assembly had a size of 1.33Gb and a contig N50 of 440Kb (Table 2).

Using this assembly, we performed two benchmarks: a first on the whole genome and a second on a specific genomic region which has been thoroughly analyzed in the Zhou et al. publication.

#### Benchmarking of polishing methods

Each polisher was launched (on a 36 cores server with 380GB of memory) iteratively six times on the input assembly to evaluate accuracy and impact of multiple rounds of correction. If needed, Illumina reads were aligned with BWA mem (v0.7.17 with the default parameters except -t 36), and the resulting bam file was sorted and indexed using Samtools(16) (v1.10 with the default parameters except -@ 36 and -m 10G). Surprisingly, HyPo never succeeded when using a genome size of 1300Mb for the correction of the *Solanum T*. genome assembly, but was able to polish the sequence when using 120Mb. The parameters used for each polisher are described in Table 1.

#### Polishing of an homozygous genome assembly - *Arabidopsis thaliana*

The Fastmer script(25) was used to generate statistics on the quality of the alignment between the polished assemblies and the reference genome (Col-0 downloaded from the TAIR website), and a quality score was calculated, for each assembly, using the following formula: *QScore = -10 * log10(1 - (matches / (matches + mismatches + insertions + deletions))*. Additionally, the accuracy of the gene content was assessed by extracting exons from the TAIR10.1 annotation (using the getfasta command from bedtools(26)) and aligning the exons onto each assembly using the Blat aligner(27). Exons that were aligned with 100% identity along their entire length were kept and unique exon names were counted to avoid multi-mapping bias.

#### Synthetic diploid sequence

The polished assemblies of the original 100Kb genomic region were aligned with the two haplotype sequences using muscle(28) (version 3.8) and each position was labeled haplotype 1 or haplotype 2 (if the base was similar to the corresponding haplotype), error (if the base was different from the two haplotypes) or equal (if the three bases were identical). For each polisher, the number of swaps between the two haplotypes and the number of errors were reported (Figure 4).

**Figure 4.**
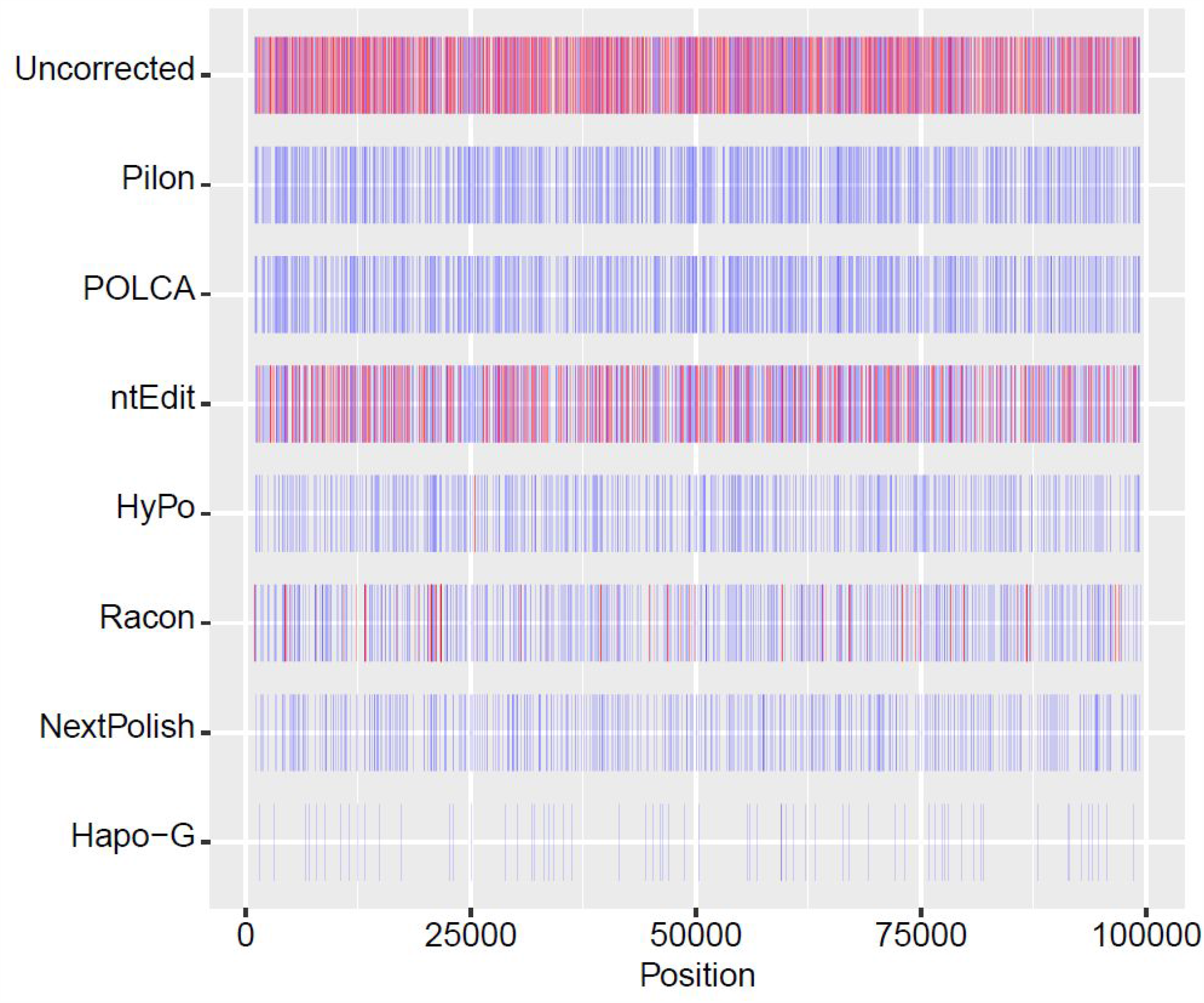
Comparison of polishing algorithms on a synthetic diploid sequence. The 100Kb sequence is represented on the x axis and each polishing tool has a dedicated track, where remaining errors are represented with red bars and switches between the two haplotypes are represented by blue bars.

#### Polishing of an heterozygous genome assembly - *Solanum tuberosum L*

The diploid and heterozygous genome of *Solanum tuberosum* was used to assess the ability of each polisher to locally preserve the haplotype phasing. Two benchmarks were performed: a first on the whole genome and a second on a specific genomic region.

Polishing algorithms were compared on their ability to phase genomic regions during the error-correction step. For that purpose, heterozygous variants were detected from the Illumina short reads, without prior assembly, using discoSNP(29) and default parameters. The sequence context (30bp on the right and left side) of each variant was extracted from the discoSNP output and only 61bp-sequence with a single variant were kept. In the discoSNP output, the detected variants were phased based on the Illumina reads (-A parameter of discoSNP), and only chains of at least three variants validated by at least 5 short reads were selected. For each heterozygous SNP, the two variants were mapped on each polished assembly and only perfect matches were kept. All reliable chains of variants were searched in the alignment results and a given chain was validated only if all its variants were found in a perfect match and on the same genomic sequence.

In addition, we focused on a 300Kb genomic region of chromosome 8, which has been described in the Figure 2b of the Zhou et al. publication. The authors illustrate a syntenic block on the two haplotypes. The coding exons of the two haplotypes were extracted and only the exons that contain at least one difference were selected and used as candidate exons. This region was assembled into four different contigs in our nanopore-based assembly, with two collapsed (contig_3372 and contig_15103) and two duplicated contigs (contig_15126 and contig_15127). The candidate exons were searched in the polished versions of these four contigs and only the perfect alignments were kept.

## RESULTS

### Polishing of an homozygous genome assembly-*Arabidopsis thaliana*

Overall, all of the polishing tools achieved very similar results in terms of quality metrics. They produced an assembly with an identity rate greater than 99.9% (Figure 3A), with the exception of Racon (99.6% after the first round of polishing). All tools, except Racon and ntEdit, produced a corrected assembly with a quality score of at least 30, right from the first round of correction (Figure S1). These results are confirmed with the number of perfectly mapped Illumina read pairs (Figure 3B), with the lowest scores obtained by Racon and ntEdit (34.1M and 41.8M respectively), while the highest number was achieved by HyPo and NextPolish (43.0M after the first round of correction). Regarding the alignment of the reference annotation, again, differences were small (less than 450 exons between Hapo-G, HyPo, NextPolish, Pilon and POLCA out of the 203,233 input exons), with ntEdit and Racon assemblies containing the lower number of exons retrieved perfectly (Figure 3C). For Hapo-G, HyPo, ntEdit, NextPolish, Pilon and POLCA, increasing the number of polishing rounds didn’t seem to have any significant impact, increasing average identity rate by less than 0.01%. Oddly, increasing the number of polishing rounds with Racon decreased the average identity rate from 99.6% for the first round to 99.1% for the sixth round. Racon’s inferior performance can be explained by the fact that it was originally designed to perform polishing using long reads. The fastest average running time was achieved by ntEdit with approximately 25 minutes for each round, while the slowest was Racon, with an average running time of 163 minutes. From the alignment-based methods, Hapo-G was the fastest with an average running time of 40 minutes (Figure 3D).

### Polishing of a synthetic sequence

Initially, the 100Kb sequence contained 861 haplotype switches and 1,877 sequencing errors. In this benchmark, we only performed one round of correction for each tool. Pilon was the only polisher to generate more haplotype switches than there were initially in the reference (881). The POLCA and ntEdit polished sequences contain more than 800 switches, while about 500 switches were still present in the HyPo, Racon and NextPolish corrected sequences. Hapo-G, the only tool dedicated to heterozygous genomes, obtained the best result with only 65 switches (Figure 4 and Table S1). In terms of remaining sequencing errors, only three corrected sequences still contain some errors, the one obtained with: HyPo (5 errors), Racon (198 errors) and ntEdit (938 errors).

### Polishing of a heterozygous genome assemblies

In the case of a more complex and heterozygous genome, the situation is different. Three methods seem to perform better than the others: Hapo-G, HyPo and NextPolish. As for simple and homozygous genomes, ntEdit and Racon obtained the worst results (Figure 5A). However the number of phased variants that could be recovered is higher in the assembly corrected with Hapo-G and this from the first round (472,534 phased variants compared to 469,441 after six rounds of NextPolish, Figure 5B). The situation is the same when looking at chains of at least three variants. Hapo-G is the only one to retrieve more than 100,000 chains whereas the second best result is obtained by NextPolish with 92,291 chains (Figure 5C). Additionally, Hapo-G is twice as fast as HyPo and seven times faster than NextPolish, the other two methods that perform well on heterozygous genomes. Furthermore, Hapo-G is the second fastest method and as previously observed ntEdit is the fastest (Figure 5D). The six rounds of polishing using Hapo-G were faster than a single round performed with NextPolish. Interestingly, we have observed three types of tools: those that take advantage of multiple rounds of correction, those for which one round seems sufficient, and those that produce inferior results by performing multiple rounds of correction. Hapo-G, HyPo, Pilon and POLCA are in the first category, ntEdit and NextPolish in the second and Racon is the only one which seems to degrade the quality of the assembly as rounds are performed (Figures 5A, 5B and 5C).

**Figure 5.**
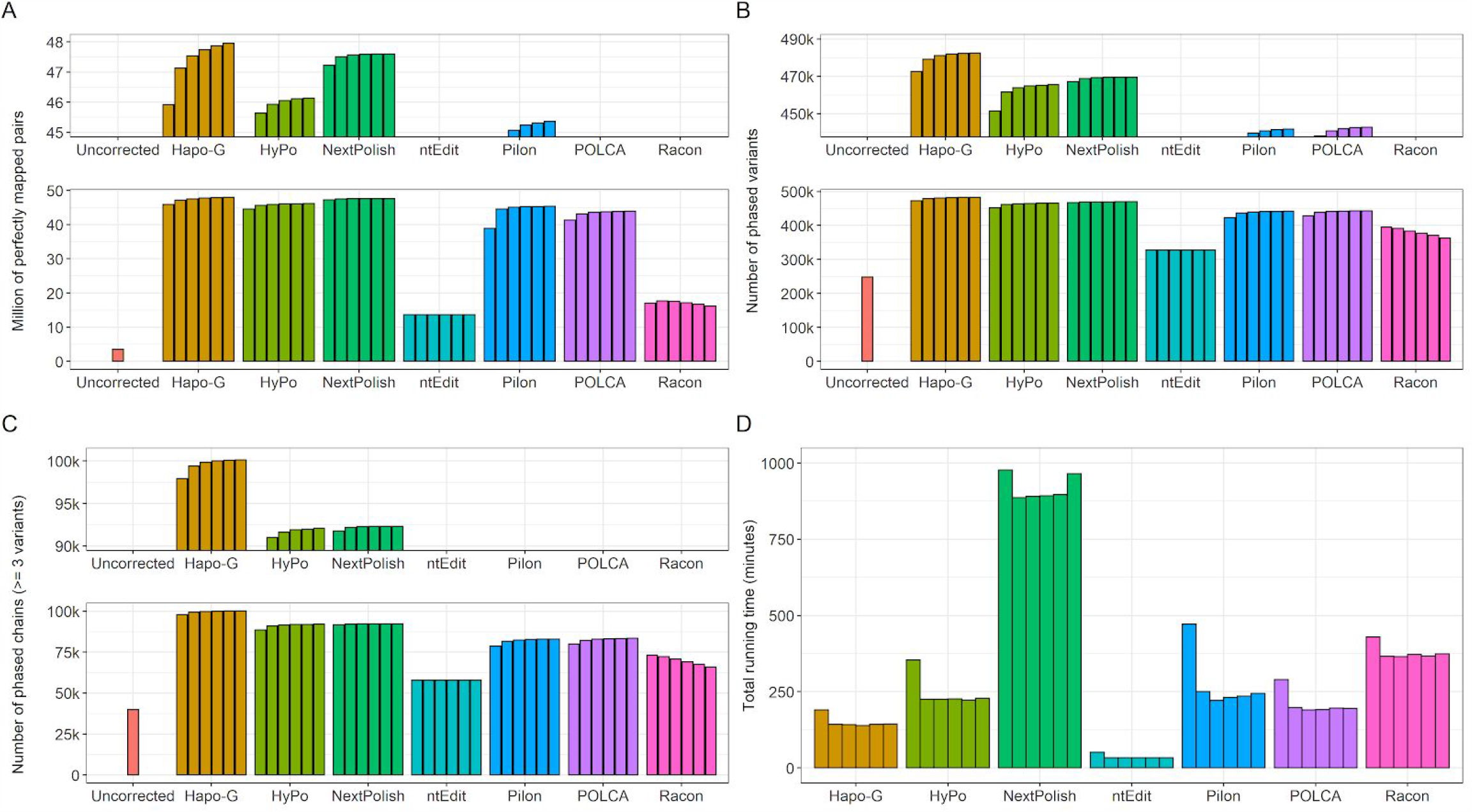
Comparison of polishing algorithms on the *Solanum tuberosum* genome assembly for each polishing round. Lower panels of A, B and C show the full distribution and the upper panels are a zoom on the higher values. **A**. Number of Illumina pairs mapped perfectly on each assembly. **B**. Number of phased variants retrieved in each assembly. **C**. Number of chains composed of more than 3 variants (and confirmed by at least 5 reads) perfectly retrieved in each assembly. **D**. Run time of each polishing tool.

In addition, we focused on the two allelic regions described in the study by Zhou et al., and counted the number of candidate exons in each assembly. In the Hapo-G corrected sequence, we recovered 86 candidate exons which represent the highest number of exons found in all assemblies, compared to 39 in the unpolished assembly. For comparison, 84 candidate exons were found in the NextPolish assembly which is the second best result (Table S2).

## DISCUSSION

In this study, we report a new software, Hapo-G, which is able to polish draft assemblies with a quality equivalent to that of existing tools on simple and homozygous genomes, while being faster, but which also improves the polishing of heterozygous genomic regions.

Nowadays, the number of polishing tools is high and although almost all are based on the same principle (mapping of short reads), their performances are different, likewise none are specialized in processing heterozygous genomes. In this study, we compared seven existing algorithms: Hapo-G, HyPo, NextPolish, ntEdit, Pilon, POLCA and Racon. We obtained very similar results on a small plant genome (*Arabidopsis thaliana*) with the exception of ntEdit which is the only tool not based on short-read alignment. In addition, we observe that the oldest and most widely used polishing tool, Pilon, is not among the tools with the best results.

We observed on a synthetic diploid sequence and on a true heterozygous genome (*Solanum tuberosum*) that the phasing of variants is of better quality when the assembly is polished with Hapo-G, leading after six rounds of correction to the higher number of paired-end reads perfectly mapped back to the assembly (47,957,836 out of 151,018,344). Only two tools, ntEdit and Hapo-G, succeeded in polishing the 1.3Gb of the *Solanum tuberosum* genome assembly in less than 3 hours on average. On this large genome, the six rounds of Hapo-G ended before the first round of NextPolish, which is the second best tool according to our benchmark and already incorporates several rounds of mapping/correction internally. Increasing the number of correction rounds is generally beneficial, except for ntEdit and NextPolish where the results are very similar from round one through sixth, and Racon where the quality of the consensus seems to deteriorate when adding new correction cycles. Based on these observations, we recommend to use Hapo-G to polish long-read assembly and eventually by performing multiple rounds of correction if possible.

In homozygous regions, although many polishers have achieved similar results, NextPolish appears to be the best performer, therefore, if possible, we suggest using NextPolish in combination with Hapo-G to achieve high quality in homozygous and heterozygous regions. In fact, homozygosity is generally not complete and heterozygous regions may remain. Interestingly, by combining NextPolish and Hapo-G, the number of perfectly mapped paired-end reads was higher than after six rounds of Hapo-G or NextPolish separately. However, a combination of any other polisher did not lead to better results in our tests (Figure S2).

## AVAILABILITY

Hapo-G is an open source software, source code, binaries as well as results of the benchmark are freely available from http://www.genoscope.cns.fr/hapog. All data, short and long reads, used in the article are available on public repositories.

## SUPPLEMENTARY DATA

All the supporting data are included in three additional files which contain a) Tables S1-2 and Figures S1-S2, b) Table S3 that contains comparative metrics on the *Arabidopsis thaliana* assembly, c) Table S14 that contains comparative metrics on the *Solanum tuberosum* assembly.

## Supporting information

Table S3: Arabidopsis results

Table S4: Solanum results

## ACKNOWLEDGMENT

The authors thank Pierre Peterlongo for his support and advice with discoSNP and his proofreading of the manuscript.

