## Supplementary Tables S1-S2 and Figures S1-S2 for "Hapo-G, Haplotype-Aware Polishing Of Genome Assemblies"

**Supplementary Table 1.** Number of haplotype switches and remaining errors in each polished synthetic sequence.

| Polisher | Number of Haplotype switches | Number of errors |
|---|---:|---:|
| Pilon | 881 | **0** |
| Uncorrected | 861 | 1877 |
| POLCA | 847 | **0** |
| ntEdit | 803 | 938 |
| HyPo | 532 | 5 |
| Racon | 520 | 198 |
| NextPolish | 438 | **0** |
| Hapo-G | **65** | **0** |

**Supplementary Table 2.** Number of candidate exons retrieved in each polished region (300Kb region from chromosome 8 of *Solanum tuberosum*).

| Polisher | Number of candidate exons (round 1, 2, 3, 4, 5 and 6) |
|---|---:|
| Hapo-G | 86,86,86,**87**,86,**87** |
| NextPolish | 84,84,84,84,84,84 |
| HyPo | 80,82,83,83,83,84 |
| Pilon | 79,84,84,85,85,85 |
| POLCA | 78,79,81,81,81,81 |
| ntEdit | 66,66,66,66,66,66 |
| Racon | 60,62,59,55,59,54 |
| Uncorrected | 40 |

**Supplementary Figure 1.** Quality score of assemblies after each round of polishing, when compared to the *Arabidopsis thaliana* reference genome. Lower panel shows the full distribution and the upper panel is a zoom on the higher values.

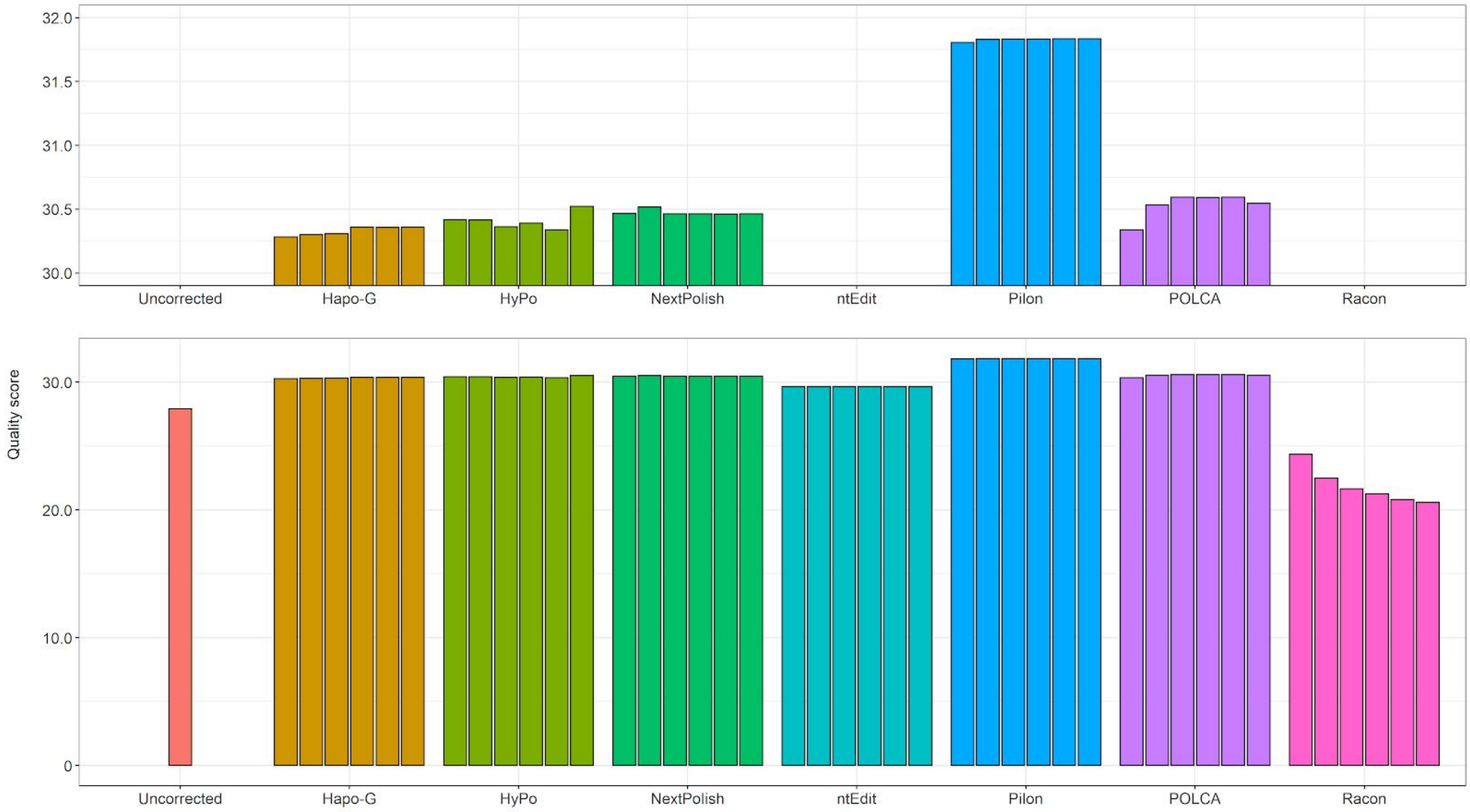

**Supplementary Figure 2.** Comparison of polishing algorithms on the *Solanum tuberosum* L. genome assembly for each polishing round. Each polisher was launched on a polished assembly obtained using Hapo-G (one cycle).

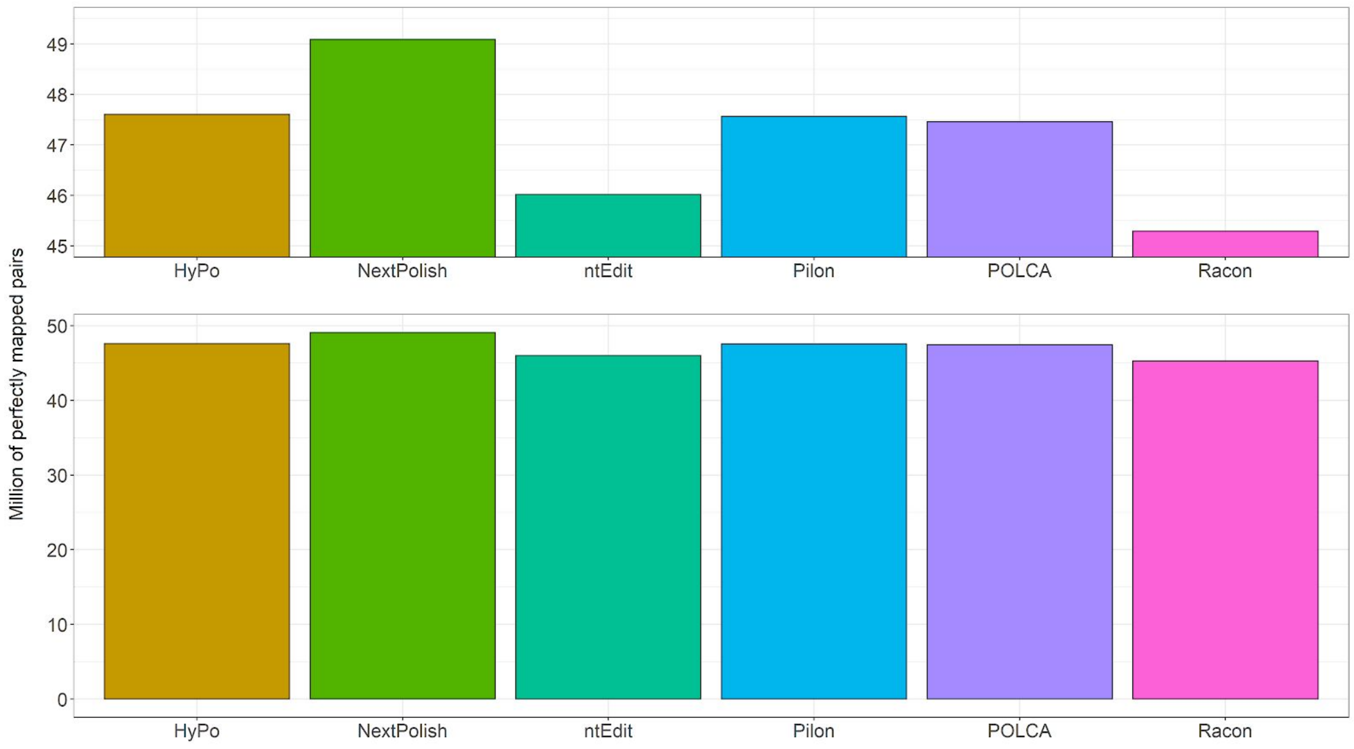
